# Comparative analyses of chromatin landscape in white adipose tissue suggest humans may have less beigeing potential than other primates

**DOI:** 10.1101/524868

**Authors:** Devjanee Swain-Lenz, Alejandro Berrio, Alexias Safi, Gregory E. Crawford, Gregory A. Wray

## Abstract

Humans carry a much larger percentage of body fat than other primates. Despite the central role of adipose tissue in metabolism, little is known about the evolution of white adipose tissue in primates. Phenotypic divergence is often caused by genetic divergence in *cis*-regulatory regions. We examined the *cis*-regulatory landscape of fat during human origins by performing comparative analyses of chromatin accessibility in human and chimpanzee adipose tissue using macaque as an outgroup. We find that many *cis*-regulatory regions that are specifically closed in humans are under positive selection, located near genes involved with lipid metabolism, and contain a short sequence motif involved in the beigeing of fat, the process in which white adipocytes are transdifferentiated into beige adipocytes. While the primary role of white adipocytes is to store lipids, beige adipocytes are thermogeneic. The collective closing of many putative regulatory regions associated with beiging of fat suggests an adaptive mechanism that increases body fat in humans.

## Introduction

Humans have a remarkable amount of body fat. While other primates have less than 9% subcutaneous fat in the wild, the derived state in healthy humans is to maintain 14-31% body fat (1, 2). Although little is known about white adipose tissue (WAT) evolution in primates, a growing body of evidence suggests that humans have uniquely adapted WAT to support the high energy needs of our brains (1, 3–9). To better understand the evolution of increased body fat in humans, a direct comparison between human and primate adipose tissue is needed.

Here we present a comparative analysis of the chromatin landscape in human and chimpanzee WAT. We mapped <u>o</u>pen <u>c</u>hromatin <u>r</u>egions (OCRs), which are highly enriched for enhancers, promoters, and other transcriptional regulatory elements. We used macaque WAT to polarize specific open chromatin changes to either the human or chimpanzee branch. We detected 3148 regions that are differentially accessible between human and chimpanzee. Notably, we find that OCRs that are more closed in humans relative to chimpanzee and macaque are enriched for conservation and are specifically near genes involved with lipid metabolism. These regions are also enriched for a sequence motif that binds a transcription factor involved in browning of fat. The data hint at a molecular mechanism driving increased WAT accumulation in humans by shutting down beigeing pathways through chromatin regulation.

## Results

### Open Chromatin Regions profiles are unique to species

We generated ATAC-seq (Assay for Transposase-Accessible Chromatin sequencing) data on white adipose samples from humans, chimpanzees and macaque (Supplemental Table 1) (10). We mapped reads from each technical replicate to the sample’s native genome assembly. For non-human primates, we only retained reads that could be reciprocally converted between hg19 and the native genome using the genome conversion tool liftOver (11). To prevent mapping biases, we performed a reciprocal liftOver from hg19 to panTro4 (chimpanzee) and back to hg19 for human samples. We called OCR peaks for each biological replicate using MACS2(12) and generated a union set of OCRs from all three species. OCRs that contained zero reads for any sample, which is an indication of mapping problems, were removed from the analysis. Our final peak set contained 160,625 OCRs (Supplemental Table 2). We used adipose ChromHMM predictions to characterize the function of OCRs (Supplemental Figure 1, Supplemental Table 3)(13). Eighty-seven percent of OCRs are located >5 Kb from the closest transcription start site, which indicates ATAC-seq can identify distal regulatory regions in WAT (Supplemental Figure 1).

To understand general patterns of OCRs, we performed principal component analysis (PCA) on normalized count data (Figure 1B). The first eigenvector explains 67% of the variance and separates macaque samples from chimpanzee and human samples. The second eigenvector explains 23% of the variance and separates human and chimpanzee samples. Technical replicates correlate highly (Pearson > 0.85) and are more similar to one another than biological replicates within a species (Supplemental Figure 1). Like most genetically driven phenotypes, OCR profiles reflect the known primate phylogeny, which indicates ATAC-seq data can be used to analyze adipose evolution in primates.

**Figure 1.**
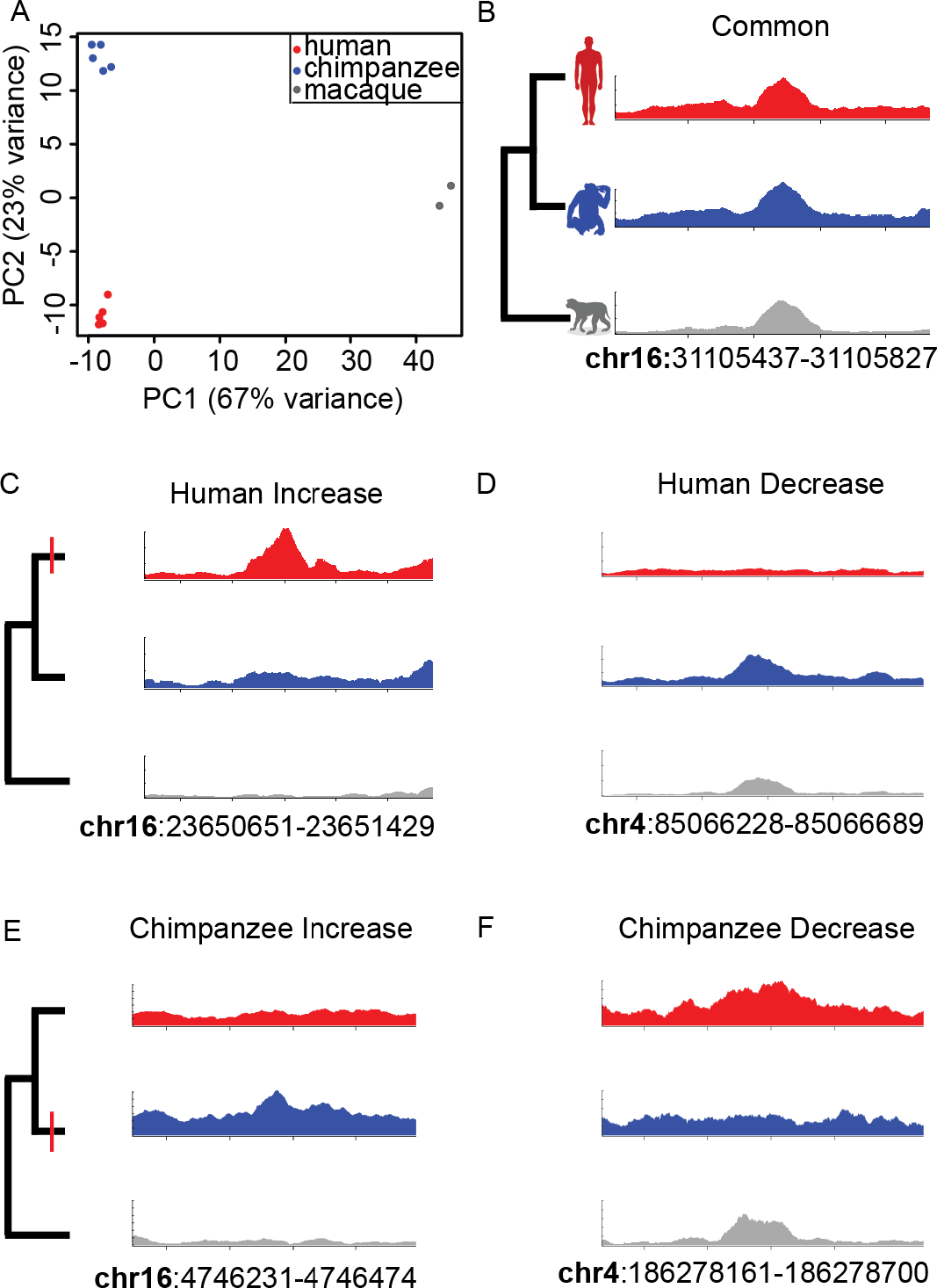
Detection of species-specific OCR state changes. (A) Principal component analysis of OCRs in human, chimpanzee and macaque adipose. Note that intra-specific variation is much smaller than inter-specific variation. A common OCR state is depicted in (B). Human-specific OCR state changes (red dash) to increased accessibility (C) and to decreased accessibility (D) from ancestral state (i.e. macaque accessibility). Chimpanzee-specific OCR state changes (red dash) to increased accessibility (E) and to decreased accessibility (F) from ancestral state. Genomic coordinates of the peak of interest are listed.

We next used DESeq2 to identify OCR regions that are quantitatively more or less accessible between species. We quantified OCR accessibility rather than simply annotate the presence or absence of a peak in a species. Since accessibility is a continuous trait, setting a threshold for presence or absence of a peak can be arbitrary and difficult to find the appropriate threshold. We also increase the number of species-specific peaks observed and increase the power for downstream analyses when we quantify OCR accessibility rather than treating accessibility as a binary trait.

Using macaque as an outgroup to assign OCR state changes to either the human or chimpanzee branch (14), we defined four groups of species-specific state changes (Figure 1, Table 1). Human increased states (n= 745) are OCRs that display similar accessibility between the chimpanzee and the ancestral state (i.e., macaque), but there is increased accessibility specifically on the human branch. Human decreased states (n= 868) consist of OCRs that display similar accessibility between chimpmanzee and the ancestral state (macacque), but there is a decreased accessibility specifically on the human branch. Chimpanzee increased (n= 1037) or decreased (n= 498) state changes are analogous to those in humans. Species-specific OCRs are increased or decreased by at least 50% in comparison to OCRs that are not classified as different between the three species.

**Table 1.**
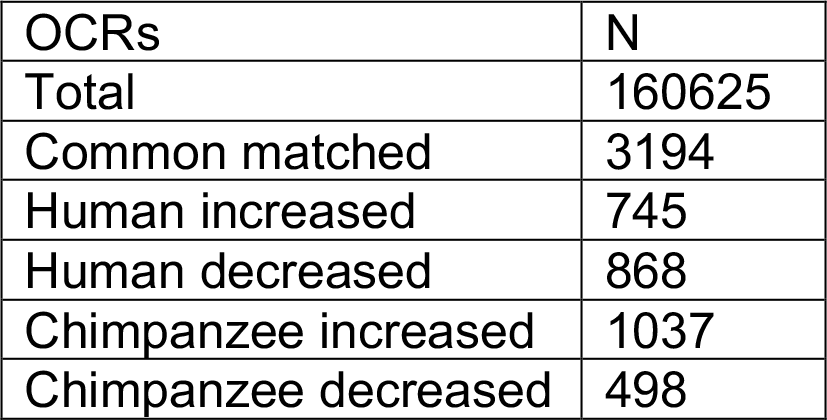
OCR groups

Our analysis resulted in 98% of OCRs as not being classified as different between humans, chimpanzees, and macaque. The complete group of nonsignificant “Common” OCRs displayed a wide range of accessibility intensities that included but does not reflect the intensities of species-specific OCRs. To ensure our downstream analyses used a control group that mirrored the intensity of the species-specific OCRs, we created a subset of matched Common OCRs that had ATAC-seq read counts between the 20-80^th^ percentiles of the species-specific ATAC-seq read counts (Supplemental Figure 1)(15).

### Species-specific OCR states correlate with *cis*-regulatory divergence

To understand the relationship between OCR state and *cis*-regulation, we assigned putative function to each OCR using publicly available human adipose ChromHMM predictions (Supplemental Table 3)(13). Approximately 13% of common OCRs are predicted to be promoters (Figure 2A). OCRs classified as being a human-decrease or chimpanzee-decrease are highly enriched for promoter regions (39.7% and 25.7% respectively, Fisher’s Exact Test, p < 0.001). In contrast, OCRs classified as a human-increase or chimpanzee-increase are significantly depleted for promoters (5.2% and 8.8% respectively, Fisher’s Exact Test, p < 0.001).

**Figure 2.**
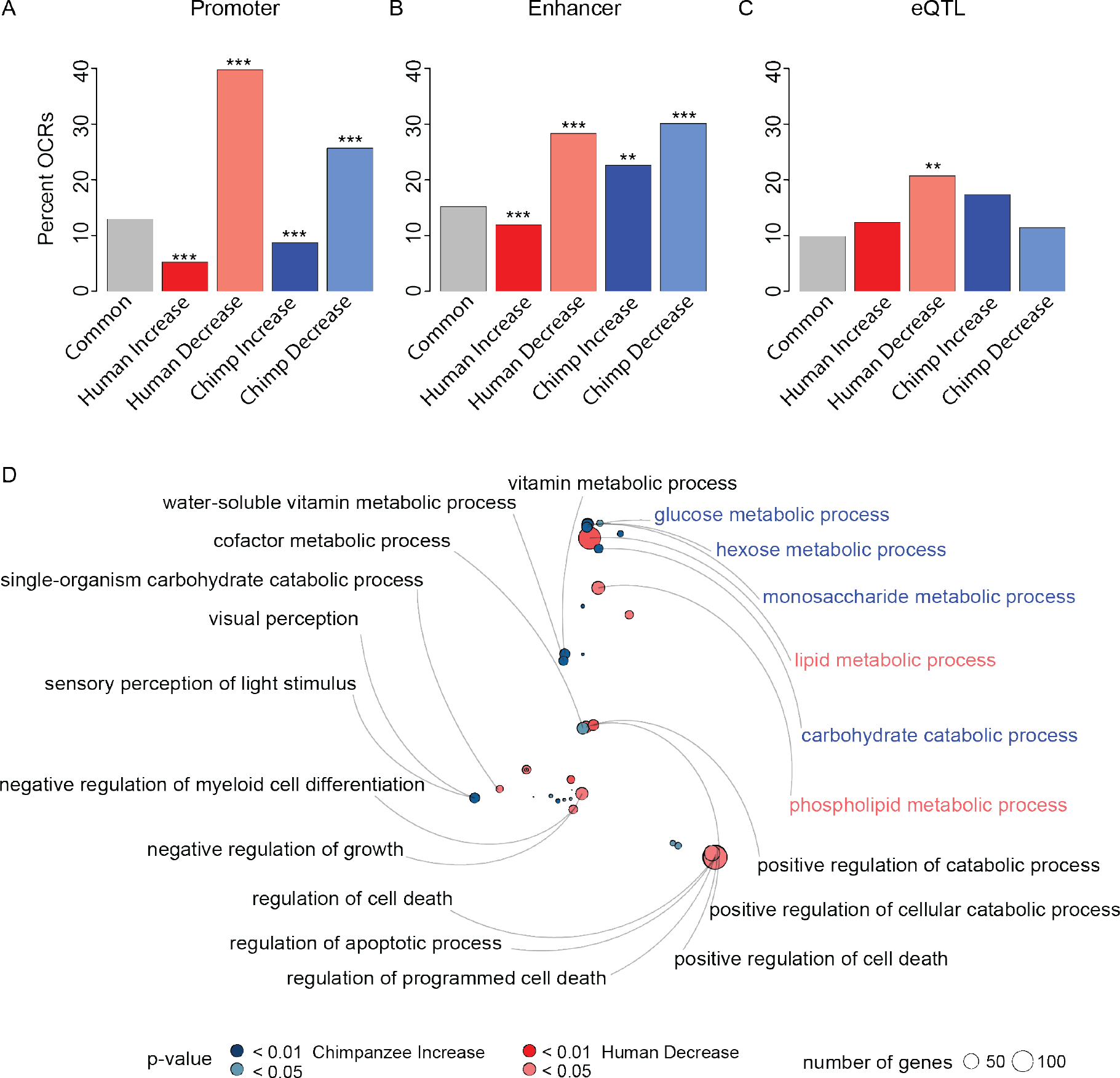
Species-specific OCR groups are enriched for *cis*-regulatory functions. Species-specific OCR groups enrichment (Fisher’s Exact, ** P <0.01, *** P <0.001) for promoters (A) enhancers (B) and adipose eQTLs (C). GREAT enrichment bubble plot (D) with labeled GO terms for bubbles containing at least 25 genes.

We next compared enhancer ChIP-seq predictions amongst OCR groups (Figure 2B). About 15% of common OCRs are predicted to be enhancers. Similar to promoters, human-decrease and chimpanzee-decrease OCRs are highly enriched for enhancers (28.3% and 30.1% respectively, Fisher’s Exact Test, FDR<0.001). Also similar to promoters, human-increase and chimpanzee-increase OCRs are not as highly enriched for enhancers. However, chimpanzee-increase OCRs displayed a higher overlap with enhancers (22.7%, Fisher’s Exact Test, p = 0.008) compared to human-increase OCRs (11.9%, Fisher’s Exact Test, p < 0.001).

These observations of promoter and enhancer enrichment and depletion reflect expected differences in the pleiotropic effects of OCR state changes in *cis*-regulatory elements. Promoters are necessary and sufficient for basal gene expression, and while enhancers can be necessary for higher expression of some genes, they are not required for low levels of expression. Furthermore, promoters tend to be pleiotropic and function in various cell types, while enhancers are mostly cell type specific(16). Finally, promoter sequence and function are more conserved than in enhancers(17). The hierarchical importance, pleiotropy, and conservation of promoters compared to enhancers implies that it is less likely to gain accessibility in promoters than in enhancers. As expected, human-decrease and chimpanzee-decrease state change are more likely to be annotated as promoter than groups with a species-increased state.

The enrichment of species-specific OCR states for *cis*-regulatory regions suggests that species-specific OCR state may be associated with functional expression changes. An association with species-specific OCRs and expression changes would support that the state changes are biologically relevant. To measure association with expression changes, we compared to known human adipose <u>e</u>xpression <u>q</u>uantitative <u>t</u>rait <u>l</u>oci (eQTL)(18).

To determine whether expression changes were enriched in species-specific OCRs, we mapped eQTL to OCRs (Figure 2C, Supplemental Table 5)(18). Interestingly, human-decrease and chimpanzee-increase OCRs are highly enriched for adipose eQTLs in comparison to common OCRs (Figure 2A, Fisher’s Exact Test, FDR = 0.002). Conversely, common, human-increase and chimpanzee-decrease OCRs are not enriched for adipose eQTLs. We note that eQTLs have thus far only been identified in humans, and thus cannot determine whether the same eQTL exists in the chimpanzee population.

Since some species-specific OCRs are enriched with eQTL, we posited that they could also be enriched for differential gene expression between human and chimpanzee. To test this, we assigned each OCR to the closest transcription start site and compared to published RNA-seq data of WAT from human and chimpanzee (Supplemental Figure and Supplemental Table 4)(19). About 5% of common OCRs are near genes associated with differential gene expression between humans and chimpanzees. Although species-specific OCRs are associated with higher levels of differential gene expression, this increase is not statistically significant.

We next asked whether species-specific OCR states were associated with biological functions. We used GREAT to perform gene ontology enrichment analyses for each OCR category (20). Similar to the eQTL analyses, Human-decrease and chimpanzee-increase OCRs are enriched for adipose-relevant gene ontology functions. In particular, they reflect the different diets of the two species: Human-decrease OCRs are located near genes associated with lipid metabolism, while chimpanzee-increase OCRs are located near genes associated with simple sugar metabolism (Figure 2E, Supplemental Tables 6-7).

To further explore the importance of species-specific OCR states with biological function, we used a branch-specific test of positive selection using the framework developed by Haygood *et al.* (9, 21). This framework compares likelihood models of neutral evolution to models of positive selection to produce a significance value associated with rate acceleration. This p-value is often correlated with the rate of evolution, ζ, which is analogous to the measure of selection in coding regions, ω. We used this framework to compute a p-value for each OCR. We compared human-branch specific selection in human-specific OCRs to that in Common OCRs and chimpanzee-branch specific selection for chimpanzee-specific OCRs to that in Common OCRs (Figure 3A and 3B). While species-specific OCRs are not enriched for more selection in comparison to Common OCRs, the strength of human-branch specific selection is significantly lower in human-specific OCRs than in Common OCRs (Figure 3A, Wilcoxon Test, Human Increase P ≪ 0.001, Human Decrease P = 0.006).

**Figure 3.**
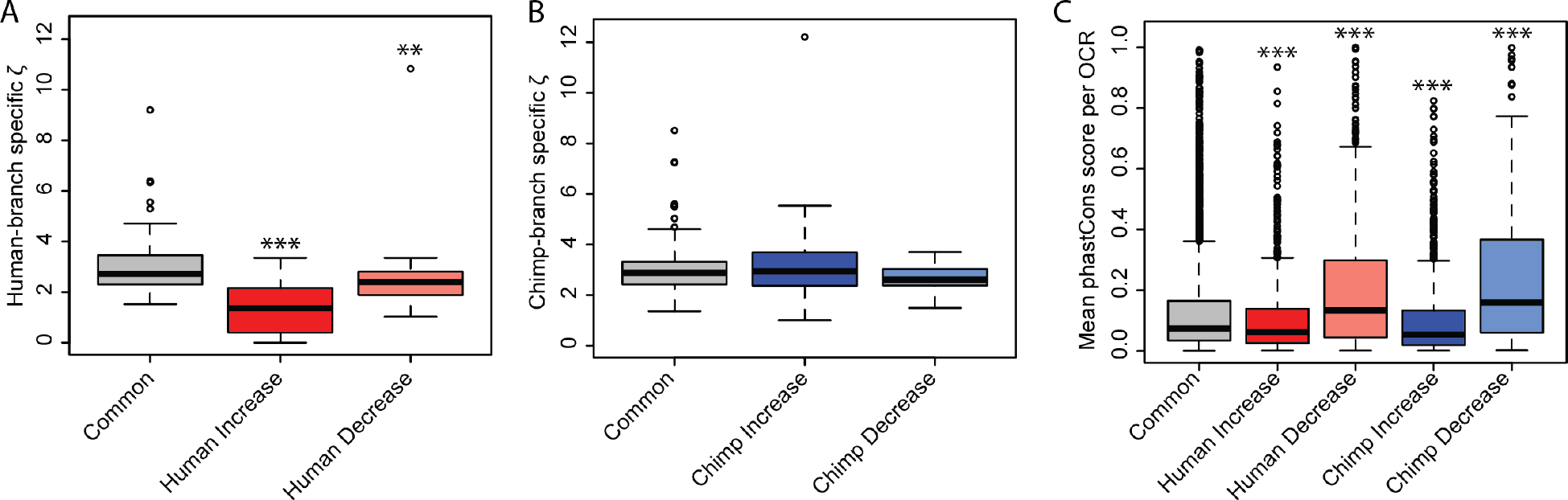
Branch-specific positive selection. Ζ for OCRs under positive selection for human-(A) and chimpanzee-(B) branch-specific selection (Wilcoxon Test ** FDR <0.01, *** P <0.001). phastCons scores (C) (Wilcoxon Test, *** P <0.001).

Species-specific OCRs under positive selection, regardless of species or state-change, are enriched for being closest to genes involved with biologically plausible functions for adipose tissue, including browning of fat, cell differentiation, leptin regulation, and obesity and related diseases (Supplemental Table 8). Although each group has equivalent amounts of human and chimpanzee branch-specific positive selection, there is little overlap in the genes that are evolving under positive selection on the human and chimpanzee branches. This indicates that humans and chimpanzees are possibly undergoing positive selection for different phenotypes.

Since the strength of ζ was low in Human OCRs, we hypothesized species-specific OCRs may display higher evolutionary conservation compared to common OCRs. To measure conservation in OCRs, we averaged the phastCons score across each OCR. Interestingly species-decreased OCRs have significantly higher mean phastCons scores than common OCRs (Wilcoxon Test, P ≪ 0.001), while species-increased OCRs have a marginally lower, albeit significant, mean phastCons score than Common OCRs (Wilcoxon Test, P ≪ 0.001). This suggests that species-decrease OCRs could be biologically relevant in other tissues, and to prevent pleiotropic effects, an OCR will be shut down rather than undergo positive selection.

### Transcription factor binding motifs characterize species-specific OCR states as being related to brown adipogenesis

Finally, we characterized categories of OCRs for enrichment of DNA sequence motifs. For this analysis, we used a machine learning algorithm in the R package gkm-SVM to test whether k-mers could predict species-specific OCR states as distinct from the rest of the genome(22). To control for local sequence features such as GC content, we created a nearest null set of common OCRs for each species-specific OCR state (i.e., for each species-specific OCR, the closest common OCR was used for a null comparison). To ensure the matched null set was representative of the rest of the genome, we measured the average performance of gkm-SVM to classify a positive set of matched null OCRs from approximately 1100 random sequences from common OCRs (not including any matched null sequences). The matched null sets are indistinguishable from the rest of the genome, which indicates that common sequence features define species-specific OCR categories (Supplemental Figures 2 and 3).

To identify sequence features that are enriched in species-specific OCR, we compared each species-specific category to its closest null set again using gkm-SVM. We measured the weights of non-redundant 6-mers, and find that each species-specific group is distinguishable from its closest null (Figure 4A and Supplemental Figure 3). Interestingly, we find a small set of 6-mers with higher weights that classify human-decrease and chimpanzee-increase OCRs, which correspond to NFI binding motifs (Supplemental Table 11). This result is intriguing since NFIA and the master adipogenesis transcription factor PPARG co-localize to regulate adipogenesis in brown adipocytes as well as in white adipocytes transdifferentiating into beige adipocytes(24, 25). Since co-localization of NFIA and PPARG motifs is correlated with an increase in brown adipocyte gene expression, we could measure how often NFIA and PPARG binding motifs occur in the same OCR (Figure 4B).

**Figure 4.**
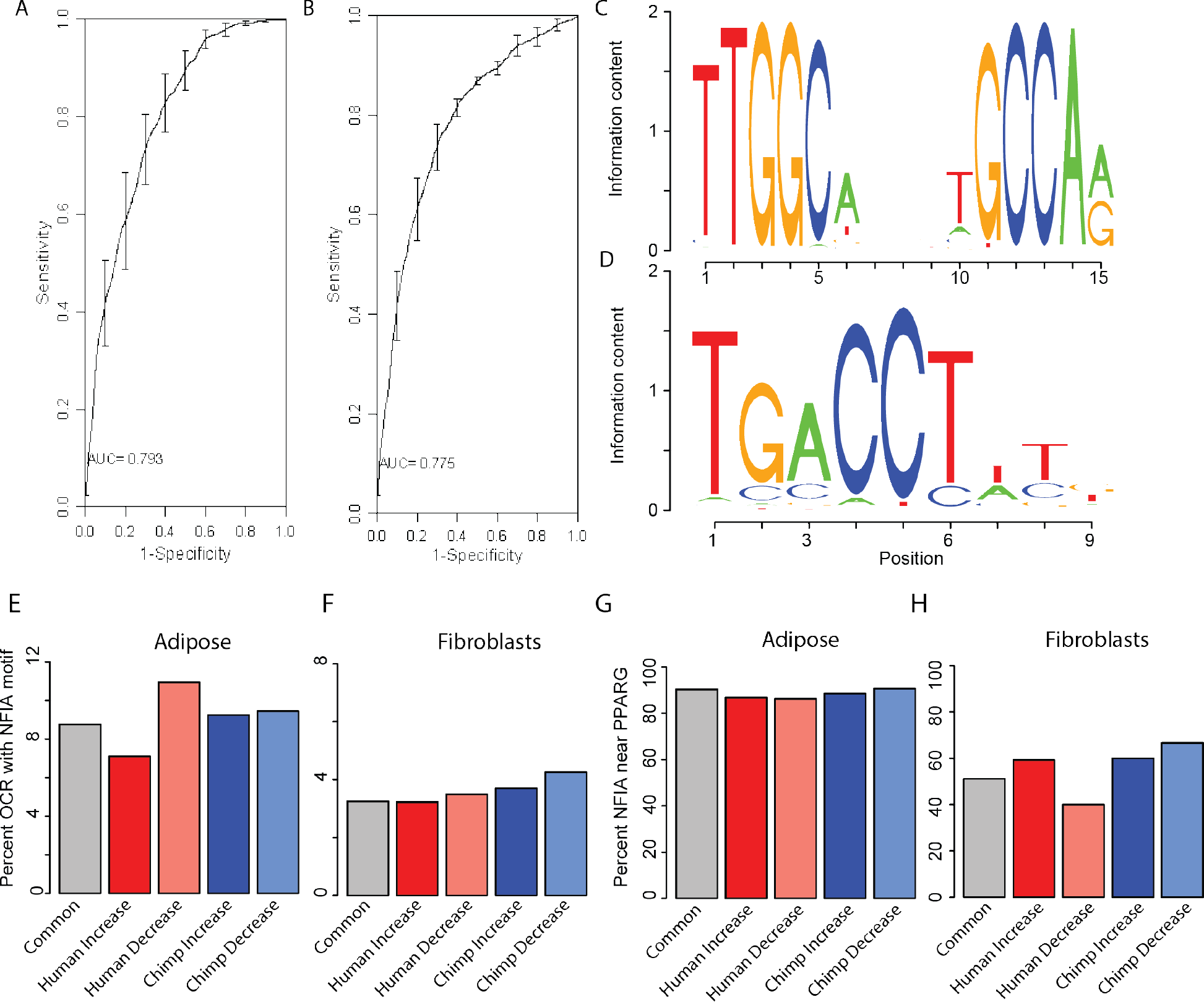
Human-decreased OCRs are associated with NFI. We used gkm-SVM to distinguish species-specific OCRs from null common OCRS. Shown are the receiver/operating curves for human-decrease (A) and chimpanzee-increase OCRs (B). We compared adipose (E) and fibroblasts (F) OCRs for NFIA (C) and PPARG (D) motifs.

To confirm that human-decrease and chimpanzee-increase OCR sequences are enriched for NFI motifs, we expanded the 6-mer motifs to the full NFIA motif and scanned all sequences for the NFIA motif(23). Human-decreased OCR have a higher percentage of OCRs (10.9%) with the longer NFIA motif than Common OCRs (8.8%), although the effect is just shy of significant enrichment (Fisher’s exact test, P=0.055). The lack of significance in species-specific NFIA enrichment could be due to the lack of confident and validated NFIA motifs. Alternatively, the NFI motif recognized could belong to three other NFI transcription factors that do not have as obvious a role in adipose as NFIA. Further a single amino acid difference between humans and chimpanzees next to the DNA-binding domain could affect binding in chimpanzees. The NFIA motif may not be the same in chimpanzee as humans, which could explain the lack of NFIA motifs in Chimpanzee Increased OCRs. To overcome this difficulty, we took advantage of NFIA’s co-localization with PPARG, the master regulator of adipogenesis.

We scanned sequences for a PPARG motif and found that over 80% of NFIA motifs occur with a PPARG motif(23). Because the PPARG motif is abundant across the genome, we wanted to ensure these observations are not an artefact and are specific to adipose OCRs. We therefore performed the same scans for NFIA and PPARG in Common and species-specific OCRs identified in a previous study in fibroblasts, which is the only study to our knowledge to also compare primate OCRs(15). We find that fibroblast OCRs have half the amount of NFIA motifs present in adipose OCRs. Additionally, only half of the fibroblast OCRs that contain NFIA motifs also contain PPARG motifs. These findings suggest that cosegregating NFIA and PPARG motifs reflect differences in biological function specific to adipose OCRs.

## Discussion

### Opposite evolutionary patterns in human and chimpanzee adipose tissue

To better understand the evolution of increased body fat in humans, we performed comparative analyses on the adipose chromatin landscape in humans, chimpanzees and macaques. Interestingly, there seem to be two modes of change in the regulatory landscape within human and chimpanzee adipose tissue. In general, human-decreased OCRs are enriched for promoters and enhancers compared to human-increased OCRs (Figures 2 and 3). Human-decreased OCRs are also more enriched for adipose eQTLs, relevant gene ontology, and NFI motifs related to adipogenesis and beiging of fat. We also find that Chimpanzee-increased OCRs are more closely associated with functional enrichment of promoters, enhancers, relevant gene ontology, and NFI than chimpanzee-decreased OCRs, Figures 2D, 2E and 3). Human-decreased OCRs are located near genes associated with lipid metabolism, while chimpanzee-decreased OCRs are located near genes associated with simple sugar metabolism. These differences in gene ontology association may reflect the differences in the diets of these two species. Taken together, these results suggest that humans shut down regions of the genome to accommodate a high fat diet while chimpanzees open regions of the genome to accommodate a high sugar diet.

### Humans may have lower beiging potential than chimpanzees

Our results further suggest a mechanism that may have contributed to the evolution of increased WAT in humans. The body contains two kinds of adipose tissue. The vast majority is white adipose tissue (WAT), which is composed primarily of white adipocytes and acts as an endocrine and lipid storage organ. In addition, the body contains brown adipose tissue (BAT), which is comprised primarily of brown adipocytes and whose main role is thermoregulation. Brown and white adipocytes differentiate from distinct mesenchymal cell populations(27, 28). White adipocytes derive from preadipocyte precursors while brown adipocytes derive from myoblasts, which can also differentiate into muscle cells. Furthermore, brown adipocytes are characterized by many small lipid droplets and a large number of mitochondria, while white adipocytes contain one large lipid droplet and fewer mitochondria.

While WAT derives from a distinct cell lineage and is predominantly made up of white adipocytes, it also contains brown-like cells, called beige or brite adipocytes. Beige adipocytes are a distinct thermogenic fat cell type from brown adipocytes; they derive from the same lineage as white adipocytes and form sporadic pockets within WAT(27–31). Beige adipogenesis is induced under a variety of conditions such as cold, caloric restriction, and exercise(27–29). Although beige adipocytes stem from the same lineage as white adipocytes, beige cells share characteristics of classical brown fat, such as higher numbers of mitochondria and smaller but more numerous lipid droplets(28). Likewise, the transcriptional profile during beige adipogenesis is unique while sharing characteristics with both white and brown adipogenesis(28).

In principle, increased WAT in humans could have evolved by shifting differentiation pathways towards white rather than beige adipocytes. Although histology on frozen adipose samples is challenging, we can still observe evidence of browning from the chromatin landscape. The NFI motif has been implicated in adipogenesis and differences in brown and white tissues (24, 25). A recent systems biology comparison of murine brown and white adipose found that open chromatin regions enriched in brown adipose contain the NFI motif and a high enrichment for GO terms involved with browning of fat (24).

Consistent with these findings, we find the NFI motif enriched in regions that are specifically closed in human WAT while open in chimpanzee WAT. Human and chimpanzee expression of *NFIA* is similar, and the NFIA motif in the observed OCRs is conserved between humans, chimpanzees and macaque. The importance of NFIA in other tissues and other developmental time points may keep the NFIA binding motif constrained. Shutting down these sequences at the chromatin level is one possible strategy to create new phenotypes without producing pleiotropic effects (Figure 3C). Taken together, these observations suggest that OCRs containing NFI motifs could be regulated epigenetically in humans to direct adipocytes to maintain a white rather than beige state

Interestingly, closing these *cis*-regulatory regions could be an adaptive response to divergence in the diets of humans and chimpanzees, as suggested by the GREAT analyses. These same regions are also enriched for positive selection during human origins, which suggests part of the collective closing of elements is related to historical adaptive pressures in humans.

## Conclusions

The data presented here point to a specific molecular mechanism in beige adipogenesis that may have contributed to the derived state of high body fat mass in humans relative to other primates. The ancestral state in non-human primates could be maintained by directing white adipose to produce more beige adipocytes. Selective pressure in humans to increase lipid storage for our metabolically demanding brains (1, 3–9) may have shaped the regulatory landscape to shut down beige pathways and redirect more adipose precursor cells towards white adipocytes. The extent to which diet and genetics play a role in accumulating white versus beige adipocytes among primate species remains unexplored. The availability of primate induced pluripotent stem cells means that future studies can begin to disentangle the effects of environment and genetic divergence during adipogenesis(32).

## Materials and Methods

### Tissue samples and ATAC-seq

The adipose tissue samples used in this study are listed in Supplemental Table 1. We obtained reproducible data from three human biological replicates (1 – 3 technical replicates each), two chimpanzee biological replicates (2 – 3 technical replicates each), and one macaque (two technical replicates). Samples were dissected from deceased individuals and sent to us as frozen samples(19). The low number of biological replicates reflects the difficulty of obtaining non-human primate tissue samples.

We homogenized 20 mg of frozen pulverized adipose tissue in nuclei isolation buffer (20 nM Tris-HCl, 50 mM EDTA, 5mM spermidine, 0.15 mM spermine, 0.1% beta meracptoethanol, 40% glycerol, 1% NP40, pH 7.5) with a dounce homogenizer. The homogenate was centrifuged at 1,100 g for 10 minutes at 4C and the pellets resuspended in resuspension buffer (10 mM Tris-HCl, 10 mM NaCl, 3 mM MgCl_2_, pH 7.4). We ran tagmentation reactions at 37C for 30 minutes, purified samples with Qiagen MinElute kits, and amplified libraries with NEB NextPCR. Duke University’s Sequencing and Genomic Technologies sequenced the libraries with the Illumina 4000 producing 150 bp paired-end reads (Supplemental Table 1).

### Data Processing, Peak Calling, and Quality Control

We used bowtie2 (33) to map reads from each technical replicate to the sample’s native genome (panTro4 for chimpanzee, hg19 for humans, and rheMac2 for macaque). For chimpanzee and macaque samples, we used reciprocal liftOver with human genome hg19 to identify homologous regions between species(11). To control for mapping biases due to disparity in genome quality, we used reciprocal liftOver with panTro4 for humans. In other words we mapped human reads to hg19, used liftOver to convert reads to the panTro4 genome, and used liftOver again to reciprocally convert reads back to the hg19 genome. Unless stated elsewhere, we used hg19 coordinates to analyze the homologous regions.

For each species, we pooled mapped reads from all technical replicates, and used MACS2 to identify open chromatin regions (OCRs)(12). We specified a shift of 100 base pairs and an extension of 200 base pairs with an FDR of 0.01. We compiled OCRs from all biological samples and removed any OCR that had 0 read counts from any technical replicate, yielding a final set of 160,625 OCRs with confident 1:1:1 homology among the three species.

### Quantitative analyses of differential OCR state

To increase the number of observed state changes in peaks, we quantified the peaks based on count data rather than presence or absence of a peak. We did not use a fold-change threshold to filter out peaks, because chromosome accessibility is a continuum and setting a threshold can be arbitrary. Additionally, noisy peaks would drop out of our differential analyses either because one or more technical replicates had 0 read counts or because a differential peak signal would not be larger than surrounding noise.

DESeq2(14) was used to normalize the count data and calculated the Pearson correlation between technical replicates. We retained replicates that correlated well with other technical or biological replicates (R>0.85) for our differential analyses. To determine whether species had an effect on OCRs accessibility, we compared a linear model with a species component (peak ~ species) to a null model (peak ~ 1) in DESeq2. We assumed the known species tree and used pairwise contrasts between species and macaque as an outgroup to determine derived OCR state changes in human and chimpanzee (FDR < 0.05). OCRs without a significant species effect (FDR > 0.05) were labeled as common OCRs. Furthermore, we wanted to ensure that the set of common peaks were similar in read intensities and size as species-specific peaks. Therefore, we created a matched common set of OCRs that fell in 20-80^th^ percentile of species-specific normalized read count and size.

### Gene expression analyses

To gain insight into *cis*-regulatory function of species-specific OCR state, we measured enrichment of OCR with eQTL and chromatin annotations (13). We used GREAT(20) to determine whether sets of OCRs possibly regulate genes that are enriched in a biological process. We used species-specific OCR states as our test regions, and the full set of OCRs for our background regions.

To associate differential gene expression with OCR state, we reanalyzed data from Babbitt *et al.(19)*. We filtered out genes with 0 reads from any biological replicate and used DESeq2 to compare a linear model with a species component (expression ~ species) to a null model (expression ~ 1). We assigned enhancers to their closest transcription start site to subset the gene expression data for each OCR group, and used Wilcoxon tests to measure differences in gene expression between OCR states.

### Selection analyses

We used the framework developed by Haygood *et al.* (9) to test for branch-specific positive selection. This framework measures the likelihood ratio of an alternative model under positive selection relative to a null model of divergence due to drift and negative selection. This test produces a p-value associated to ζ, that is analogous to ω, in which ζ< 1 is indicative of a region under negative selection; ζ= 1 is indicative of region under neutral evolution; and ζ > 1 is indicative of a region of positive selection. We compared selection of species-specific OCR states to a set of genomic regions that are predicted to be non-functional based on ChromHMM annotations(13).

### Motif analyses

To determine if OCR sequences could be differentiated from the rest of the genome, we used the default settings of the machine learning R package, gkm-SVM(22). We calculated the average performance of 100 simulations for each OCR set, using a negative group of 1100 random sequences from the total peak set. We used the default settings of gkm-SVM to predict species-specific OCR sequences from matched null OCR sequences, which consisted of the closest common OCR to a species-specific null. The match null set controls for local genomic features such as GC content. We used TOMTOM from MEME Suite(34) to identify transcription factor candidates that bind to predicted motifs from gkm-SVM. We used the R package JASPAR TFBSTools (35) to scan sequences for the NFIA (M3607_1.02) and PPARG (M6434_1.02) motifs from CIS-BP Database(Supplemental Table 11) (23).

## Acknowledgements

We thank Sasha Makahon-Moore for her bubbleplot R code, and members of the Wray lab and Raluca Gordân for helpful discussions.

## Data reporting

Raw fastq files can be found at Gene Expression Omnibus (GEO). All processed data can be found in the Supplemental Tables.

## Accession Numbers

Raw fastq files are located at GEO with accession numbers GSM3494237 - GSM3494249.

