## Supplemental Tables for "Comparative analyses of chromatin landscape in white adipose tissue suggest humans may have less beigeing potential than other primates"

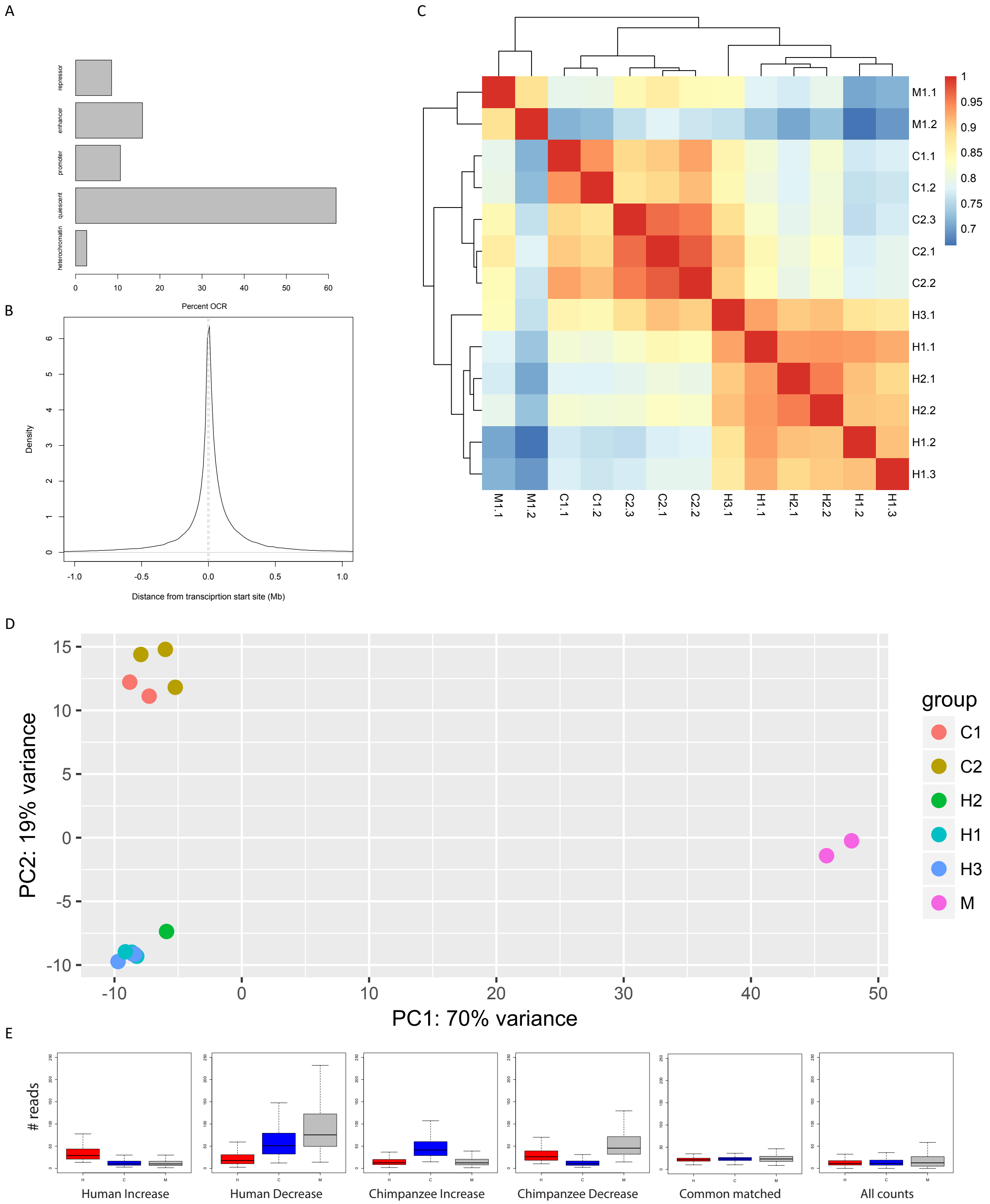


**Supplemental Figure 1. Open Chromatin Regions (OCRs) in primate adipose.** Percent OCRs annotated as functional genomic regions (A). Density plot of OCR distance from nearest transcription start site (B). Vertical gray lines are at -5 and 5 kb. Heatmap of all replicates (C). Biological replicates are H1 (Human 1104), H2 (Human 1442) , H3 (Human 602), C1 (Chimpanzee 4x327), C2 (Chimpanzee 4x519), M (Macaque) . Technical replicates are denoted by .X where X is the number of the technical replicate. PCA of OCR height for technical replicates (D). Read counts for all human (red), chimpanzee (blue) and macaque (gray) samples (E).

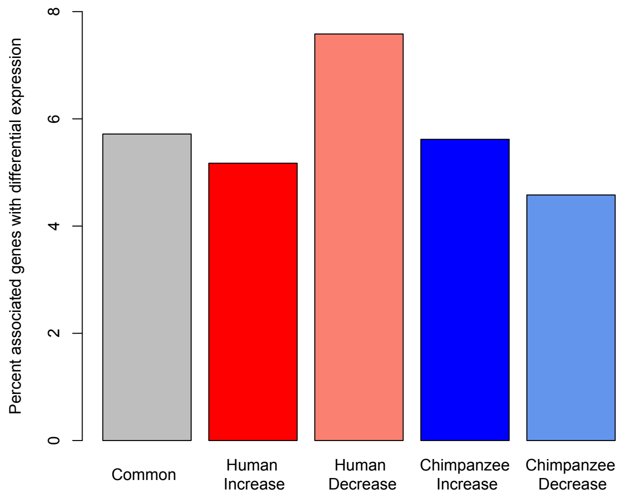


**Supplemental Figure 2.** Percent associated genes with differential expression. OCR were assigned to the closest gene.

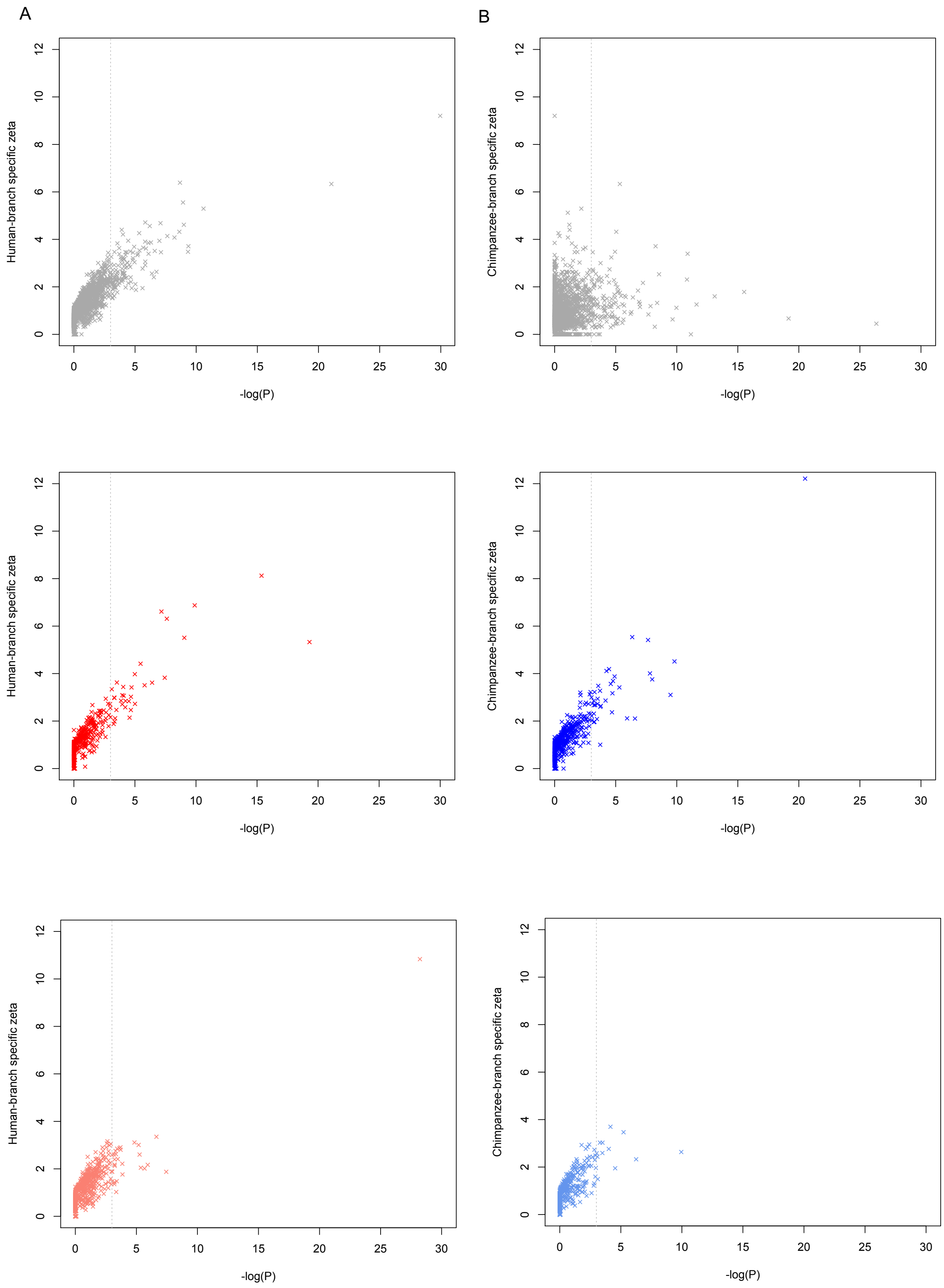


**Supplemental Figure 3.** Human- (A) and Chimpanzee- (B) branch specific selection for each OCR belonging to common (top panel), high (middle panel), and low (bottom panel) groups. The vertical gray line depicts significance ($P < 0.05$).

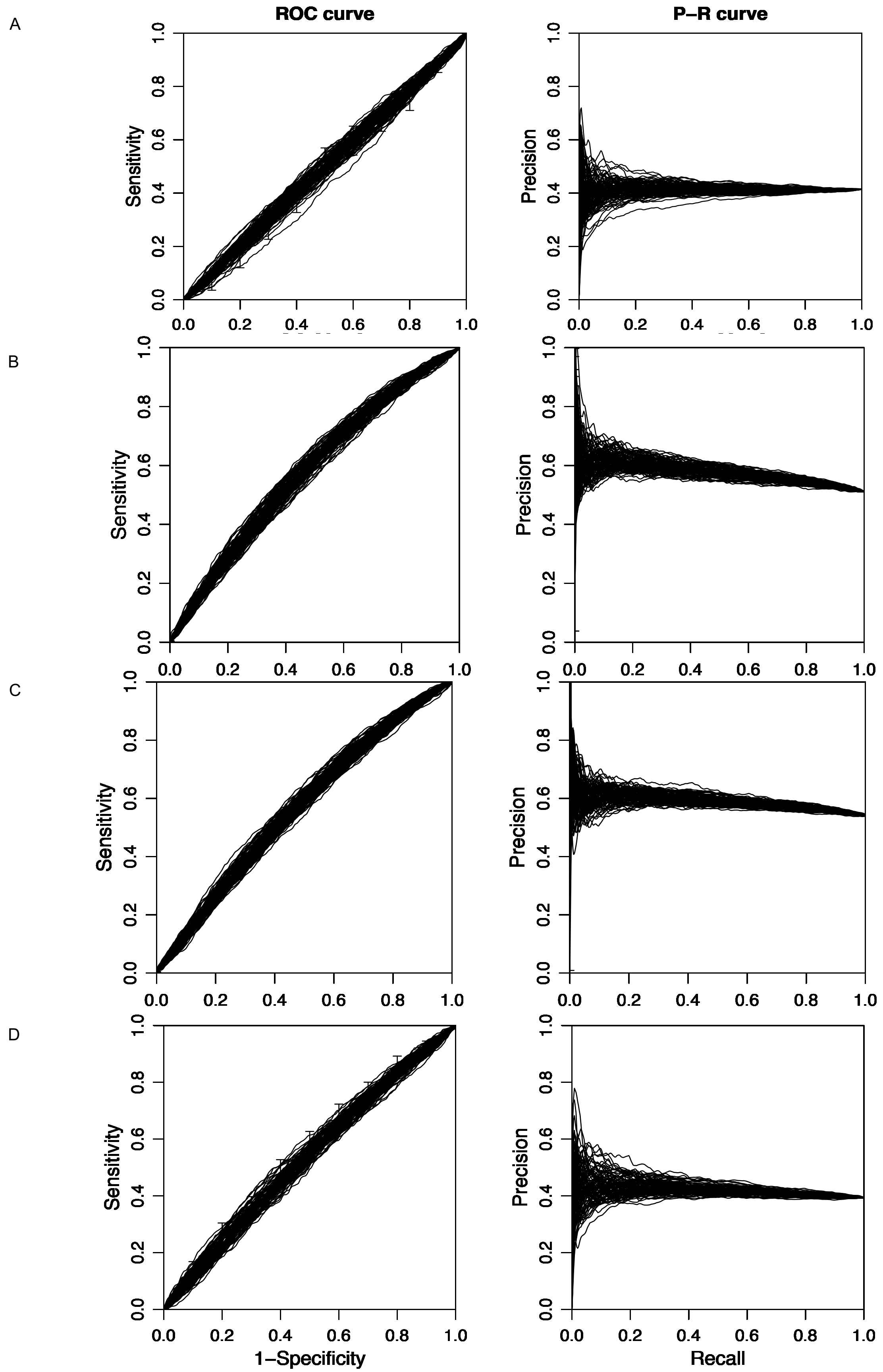
A
B
C
D
ROC curve
P−R curve
Sensitivity
Precision
1−Specificity
Recall
0.0
0.2
0.4
0.6
0.8
1.0

**Supplemental Figure 4. Comparison of matched null groups to randomized genomic sequences.** We used gkm-SVM to distinguish the matched null of human low (A), human high (B), chimpanzee low (C), and chimpanzee high (D) OCR groups from 1100 random sequences from the genome. We plotted the receiver/operating (left) and precision/recall (right) curves. Each curve is a separate set of 1100 sequences for a total of 100 sets of comparisons.

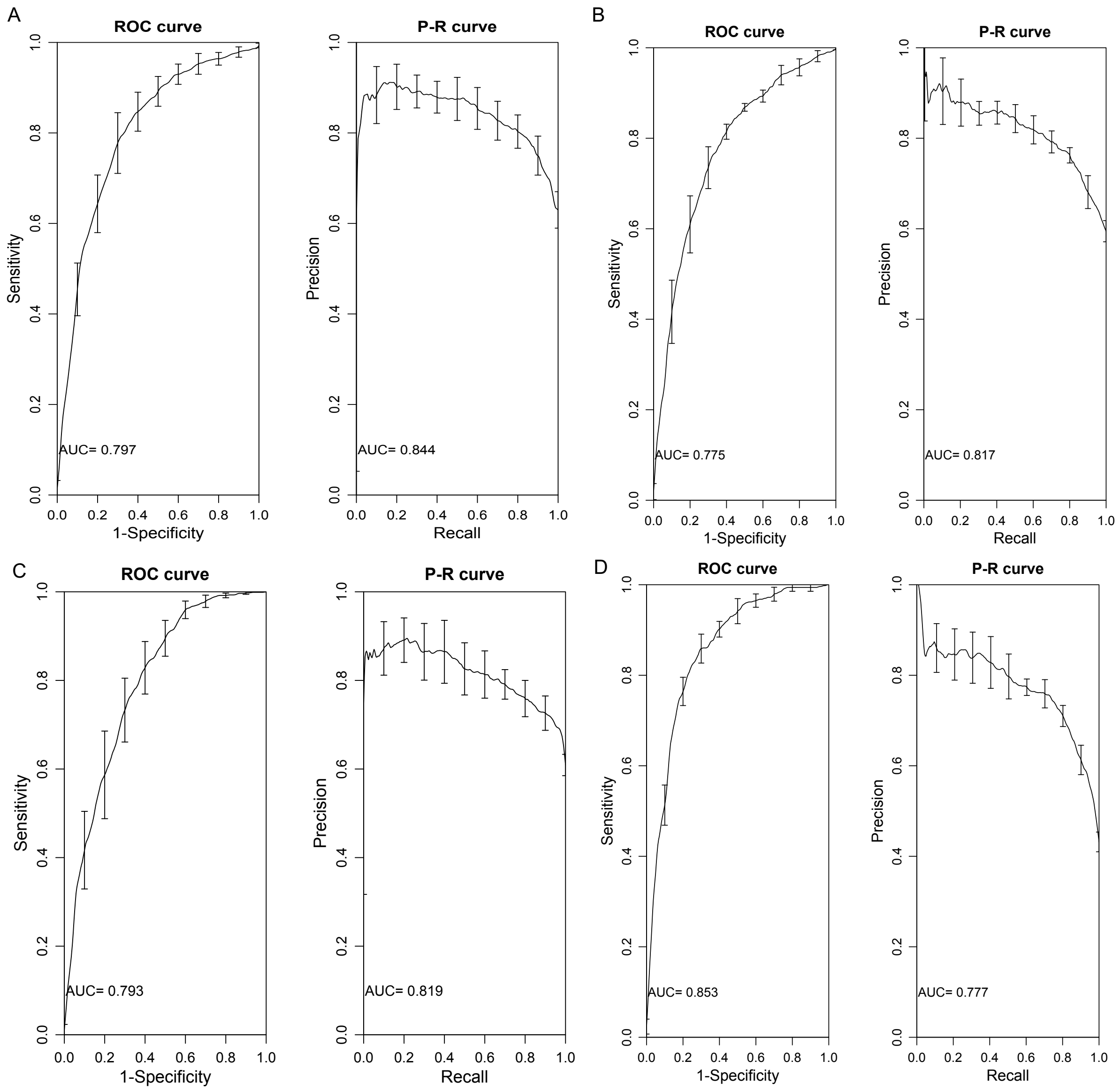


**Supplemental Figure 5.** Receiver operator characteristic (ROC) and precision recall (P-R) curves for species-specific OCR group compared to a null common group. Human High (A) Human Low (B) Chimpanzee High (C) Chimpanzee Low (D).
